# Epicardial HDAC3 promotes myocardial growth through a novel microRNA pathway

**DOI:** 10.1101/2021.09.16.460538

**Authors:** Jihyun Jang, Guang Song, Qinshan Li, Xiaosu Song, Chenleng Cai, Sunjay Kaushal, Deqiang Li

## Abstract

**Rational:** Establishment of the myocardial wall requires proper growth cues from nonmyocardial tissues. During heart development, the epicardium and epicardium-derived cells (EPDCs) instruct myocardial growth by secreting essential factors including fibroblast growth factor 9 (FGF9) and insulin-like growth factor 2 (IGF2). However, it is poorly understood how the epicardial secreted factors are regulated, in particular by chromatin modifications for myocardial formation.

**Objective:** To understand whether and how histone deacetylase 3 (HDAC3) in the developing epicardium regulates myocardial growth.

**Methods and Results:** We deleted *Hdac3* in the developing murine epicardium and mutant hearts showed ventricular myocardial wall hypoplasia with reduction of EPDCs. The cultured embryonic cardiomyocytes with supernatants from *Hdac3* knockout (KO) mouse epicardial cells (MECs) also showed decreased proliferation. Genome-wide transcriptomic analysis revealed that *Fgf9* and *Igf2* were significantly down-regulated in *Hdac3* KO MECs. We further found that *Fgf9* and *Igf2* expression is dependent on HDAC3 deacetylase activity. The supplementation of FGF9 or IGF2 can rescue the myocardial proliferation defects treated by *Hdac3* KO supernatant. Mechanistically, we identified that microRNA (miR)-322 and miR-503 were upregulated in *Hdac3* KO MECs and *Hdac3* epicardial KO hearts. Overexpression of miR-322 or miR-503 repressed FGF9 and IGF2 expression, while knockdown of miR-322 or miR-503 restored FGF9 and IGF2 expression in *Hdac3* KO MECs.

**Conclusions:** Our findings reveal a critical signaling pathway in which epicardial HDAC3 promotes compact myocardial growth by stimulating FGF9 and IGF2 through repressing miR-322/miR-503, providing novel insights in elucidating etiology of congenital heart defects, and conceptual strategies to promote myocardial regeneration.

## Introduction

Congenital heart disease (CHD) is still the most common birth defect worldwide^1^. Myogenic defects are often associated with many forms of CHD. In the past, most research has focused on identifying the intrinsic factors in the myocardium to understand the potential causes, whereas the contributions from nonmyocytes (through intercellular communications or regulations between cardiomyocytes and nonmyocytes) have been largely overlooked^2-4^. The nonmyocyte compartments such as the epicardium and epicardium-derived cells (EPDCs) are capable of regulating the development of adjacent tissues such as compact myocardium via paracrine signaling crosstalk^5, 6^. Thus, it is possible that the disrupted signaling communications other than myocardium itself account for the myocardial malformations. Further, many of these paracrine signaling pathways are reinvested during heart repair and/or regeneration. For instance, the epicardium acts as a pivotal hub for mediating heart regeneration^5, 7-9^. Studies in both adult zebrafish hearts and neonatal mouse hearts have revealed that the epicardium plays a critical role for heart regeneration by providing paracrine growth factors, such as fibroblast growth factors (FGFs) and insulin-like growth factors (IGFs) to proliferating cardiomyocytes^10-13^. Understanding how these paracrine signals are regulated may provide important insights for both the pathogenesis of CHD and novel strategies for promoting heart regeneration.

The epicardium is composed of a single layer of mesothelial cells. It covers the outermost layer of the heart. The epicardium originates from a cluster of progenitor cells, which are located at the venous pole of the developing heart, known as the proepicardium (PE)^14^. The PE is initiated around embryonic day (E) 8.5 in mice^14, 15^. The EPDCs emancipate from the epicardium through an epithelial-to-mesenchymal transition (EMT) event and give rise to several cardiac cell lineages including interstitial cardiac fibroblasts and coronary smooth muscle cells, which constitute cardiac stroma and provide oxygen and nutrients to heart muscle^16-19^. During heart development, the epicardium also plays an important role by nurturing the underlying myocardium through secreting paracrine trophic factors such as FGF9 and IGF2^4-7, 20-26^.

FGF9 belongs to the FGF super family^27^. The binding of FGF9 to FGF receptors (FGFRs) triggers phosphorylation of FGFRs, and then subsequently activates the PI3K/AKT pathway and the MEK/ERK signaling cascade to drive cell proliferation and tissue morphogenesis^27, 28^. Either global deletion of FGF9 or conditional knockout of FGFR1/2 in the myocardium leads to ventricular hypoplasia^23^, suggesting that FGF9 and its downstream signaling is important for myocardial growth. IGF2 is another major paracrine growth signal released from epicardial cells and converges to the same downstream AKT and ERK signaling axis^5, 29^. Conditional knockout of either IGF2 in the epicardium or its major receptor, IGFR1 in cardiac progenitors (Nkx2-5+) resulted in reduced cardiomyocyte proliferation and ventricular wall hypoplasia^24-26^. In contrast, conditional deletion of IGF2 in the endocardium or myocardium did not give rise to any apparent cardiac phenotypes^26^. In the developing heart, FGF9 and IGF2 are mainly secreted by the epicardium and its EPDCs, with minor contributions from cardiac endothelial cells^5, 20^. Most previous research has focused on understanding the function and/or the downstream signaling of these growth factors^27, 29^. How the expression of these growth factors in the developing epicardium is regulated is still poorly understood.

Gene transcription can be heavily influenced by chromatin*’*s accessibility, which can be regulated by post-translational modifications of histones, including acetylation/ deacetylation, methylation/demethylation, phosphorylation/dephosphorylation and ubiquitination/deubiquitination^30, 31^. By switching between any of those two states, the associated genes can be dynamically programmed to be transcriptionally active or repressed. Histone deacetylase 3 (HDAC3), a member of the class I HDAC family that catalyzes the removal of acetyl groups from lysine residues in histone tails, has been implicated in many biological processes by modulating gene expression^32^. Mesodermal or global knockout of *Hdac3* resulted in myogenic defects and early embryonic lethality^33, 34^. Interestingly, ablation of *Hdac3* specifically in the myocardium does not give rise to cardiac morphological phenotypes during early heart development, but rather compromises cardiac function at later postnatal stages^34, 35^. These findings suggest that the function of HDAC3 in nonmyocyte compartments may be critical for early myocardial development.

MicroRNAs (miRs) are small non-coding RNAs (about 22 nucleotides in length) that post-transcriptionally regulate gene expression, and are greatly implicated in heart development, disease, and regeneration^36, 37^. MiRs can modulate the expression of a wide range of genes including growth factors. For instance, the expression of FGF2 and IGF2 in neural stem cells, cancer, or skeletal muscle is subject to the regulation by miR-1275, miR-483, and others^38-41^. It is not clear whether and how the expression of FGF9 and IGF2 is epigenetically (e.g., via HDACs or miRs) regulated in the developing epicardium.

In the present study, we investigated the role of HDAC3 in the epicardium during early heart development.

## Methods

### Mice

*Hdac3*^*flox/+*^, *Tnnt2*^*nGFP/+*^, and *Wt1*^*CreERT2/+*^ mice were previously described^17, 42-44^. *Wt1*^*CreERT2/+*^ (Stock #010912) and *R26*^*eYFP*^ mice (Stock #006148) were purchased from The Jackson Laboratory (Bar Harbor, ME). We got *Hdac3*^*flox/+*^ mice from Dr. Mitchell Lazar at the University of Pennsylvania. *Hdac3*^*flox/+*^ mice (Stock #024119) are also available at The Jackson Laboratory. All animal protocols were approved by the University of Maryland Baltimore Institutional Animal Care and Use Committee.

### Administration of tamoxifen and 5-bromo-2’-deoxyuridine (BrdU) in vivo

Tamoxifen (Sigma, Saint Louis, MO) was dissolved in corn oil and intraperitoneally (IP) given to pregnant mice (150 mg/kg body weight) at E8.5 by gavage. BrdU (Sigma) was dissolved in phosphate-buffered saline (PBS) and given IP to pregnant mice (100 mg/kg body weight) at E11.5 and E12.5 (one dose per day).

### Embryonic cardiomyocyte culture and proliferation assessment

E13.5 ventricular cardiomyocytes were isolated from dissected embryonic hearts and cultured as previously described^45^. Briefly, hearts from *Tnnt2*^*nGFP/+*^ embryos were collected at E13.5 and digested in Trypsin solution (10 mM HEPES, 0.5 mM EDTA, 0.5% Trypsin in 1X HBSS solution) at 4°C for overnight on a rocker. On the following day, cells were spun down and dissociated in dissociation buffer (10% horse serum [w/v], 5% FBS, 10 mM HEPES, and 1X HBSS solution). After centrifugation, cell pellets were resuspended in rinse buffer (10% horse serum, 5% FBS, 1% P/S, and Ca^2+^ free DMEM) and filtered through 40 um cell strainers. After centrifugation, cells were resuspended in cultured media (Opti-MEM, 10% horse serum, 5% FBS, and 0.1% P/S) and cultured at 37°C in a humidified incubator with 5% CO2. To eliminate adherent nonmyocytes, floating cells were replated on laminin-coated tissue culture dishes/chamber slides after one-hour culture. To assess cell proliferation, cardiomyocytes were fixed with 4% paraformaldehyde and stained with p-H3 antibody (Supplemental Table 1) followed by Alexa 594-conjugated secondary antibody (Thermo Fisher Scientific, Waltham, MA). The immunostaining was visualized and imaged on a Leica DM6 fluorescence microscope (Wetzlar, Germany). The percentages of p-H3+GFP+/GFP+ or total number of GFP+ cardiomyocytes were quantified using ImageJ software.

### Histology and immunohistochemistry

All embryo specimens were fixed in 4% paraformaldehyde overnight, dehydrated through an ethanol series, paraffin embedded, and sectioned (6-7 μm). Primary antibodies (Supplemental Table 1) were incubated at 4°C overnight and secondary antibodies (Alexa Fluo 488, 555, or 647; Thermo Fisher Scientific) were incubated at room temperature for 1 hour. The stained slides were imaged on a Leica DM6 fluorescence microscope. We quantified the number of p-H3+ cells, BrdU+ cells, TUNEL+ cells EPDCs (eYFP+), and the staining intensity of FGF9 and IGF2 by using ImageJ software.

### Cell culture, transient transfection, lentiviral infection, and luciferase assay

Mouse embryonic epicardial cells (MECs) originally generated by Dr. Sucov*’*s group^24^ were purchased from Millipore (Burlington, MA, Catalogue #SCC187). MECs were cultured in 10% fetal bovine serum (FBS) supplemented DMEM at 37°C in a humidified incubator with 5% CO2. To generate stable *Hdac3* KO MECs, we cloned *Hdac3* sgRNA into lentiCRISPR v2 vector (Addgene #52961; Watertown, MA). sgRNA primers (targeting on *Hdac3* Exon2) were 5*’*-CACCGCATAGCCTAGTCCTGCATTA-3*’* (forward) and 5*’*-AAACTAATGCAGGA CTAGGCTATGC-3*’* (reverse). Seventy percent confluent Lenti-X™ 293T cells (Takara Bio USA, Inc., Mountain View, CA) were transfected with *Hdac3* lentiviral plasmids (*Hdac3* KO or empty vector control [EV]), PsPAX2 (Addgene #12260), and PMD2G (Addgene #12259). Seventy-two hours after transfection, supernatants containing lentivirus were collected and filtered through a 40-μm cell strainer. For *Hdac3* rescue experiments in MECs, plasmids expressing human *HDAC3* and *HDAC3*^*Y298H*^ have been previously described^46^. miR-322, miR-503 mimics, and scramble control mimics were custom-ordered from Sigma. miR-322, miR-503 mimics, or control mimics (30 pmol) were transfected to MECs using Lipofectamine RNAiMAX (Thermo Fisher Scientific). miRZip anti-miR precursor constructs for miR-322 and miR-503 cluster (miRZIP-322 and miRZip-503) and pGreenPuro Scramble Hairpin control construct were purchased from System Biosciences (Palo Alto, CA), and lentiviruses were generated according to the manufacturer*’*s protocol. MECs were cotransfected with SV40-renilla and pGL4.10-miR-322/miR-503 promoter using lipofectamine 2000. After collecting the cell lysates, firefly and renilla luciferase activity were measured by VICTOR X3 Multilabel plate reader (PerkinElmer, Waltham, MA) followed by Dual-Luciferase® Reporter Assay System (Promega, Madison, WI; Catalogue #E1910).

### RNA isolation, quantitative real-time PCR (qRT-PCR), and bulk RNA-Seq

MECs (90% confluent) and E13.5 hearts were used for RNA isolation. E13.5 hearts were microdissected in cold PBS and snap frozen in liquid nitrogen. The RNeasy Plus Mini Kit (QIAGEN, Germantown, MD) was used to extract total RNA and cDNA was generated with the Superscript III kit (Thermo Fisher Scientific). To detect mRNA, SYBR Green quantitative real-time polymerase chain reaction (qRT-PCR) was performed. PCR primers for genes of interest are listed in Supplemental Table 2. To perform mature microRNA expression analysis, purified total RNA was converted to cDNA using TaqMan™ MicroRNA Reverse Transcription Kit (Thermo Fisher Scientific, Catalogue #4366596). qRT-PCR reactions were run on StepOne Plus Real-Time PCR System (Applied Biosystems, Waltham, MA). The probe sequences for mmu-miR-322-5p, mmu-miR-503-5p, and U6 snRNA were purchased from Thermo Fisher Scientific. For bulk RNA and miRNA-Sequencing, samples were prepared following the provider*’*s guidelines (Novogene Corporation Inc, Sacramento, CA) and sequenced on an Illumina NextSeq500 (San Diego, CA). Sequencing reads were aligned to the UCSC mm10 reference genome using tophat2 and bowtie2 in R^47, 48^. Differential expression of transcripts was calculated using the cufflinks suite in R analyses^49-51^.

### Chromatin immunoprecipitation assay

MECs were crosslinked for 8 min at room temperature by adding 1% final volumes of fresh 16% formaldehyde. The crosslink was then quenched by adding 1/10 volumes of 2.5 M glycine to cells and incubating at room temperature for 5 min. Nuclei were prepared from cells according to Covaris truChIP™ Chromatin shearing kit protocol and sonicated to fragments (average fragment length: 300–500 bp) using M220 focused Ultrasonicator (Covaris, Wolburn, MA). Chromatin (10 uL) was saved for whole cell extract input. anti-H3K27Ac antibody or rabbit IgG antibody was conjugated to the Dynabeads™ protein G beads (Thermo Fisher Scientific). Chromatins (15 ug) were added to the bead solution and incubated overnight at 4ºC on a rotator. Beads were then collected and washed 4 times in wash buffer (20 mM HEPES pH 7.9, 150 mM NaCl, 1.5 mM MgCl_2_, 0.2 mM EDTA, 0.2% Triton X-100, and 10% Glycerol). After removing the wash buffer, crosslinking was reversed at 65°C for 2 hours in proteinase K buffer (20 mM Tris-HCl, pH 7.5, 5 mM EDTA, 50 mM NaCl, 1% SDS, 20 mM sodium butyrate, 200 μg/ml proteinase K). DNA was purified with phenol/chloroform/isoamyl alcohol. Using purified precipitated DNA, enrichment of the target sequences was measured by chromatin immunoprecipitation (ChIP)-qPCR using primers (Supplemental Table 3) designed against the murine miR-322/miR-503 promoter regions (−1.7kb to +1bp).

### Western blotting

Cell or tissue lysates were prepared in lysis buffer (20 mm Tris-HCl [pH 7.5], 15 mm NaCl, 1 mm Na_2_EDTA, 1 mm EGTA, 1% Triton X-100, 1 μg/ml leupeptin, 2.5 mm sodium pyrophosphate, 1 mm Na_3_VO_4_, and 1 mm β-glycerophosphate) with protease inhibitor cocktail (Roche, Indianapolis, IN) and 1 mM phenylmethylsulfonyl fluoride. Protein samples were resolved on 4–12% SDS-PAGE acrylamide gel before transferring to polyvinylidene fluoride membranes. Primary antibodies (Supplemental Table 1) were visualized by chemiluminescence using HRP-conjugated secondary antibodies. The density of protein bands was quantified by ImageJ software.

### Statistical analysis

Data are presented as mean *±* SEM. Statistical significance was determined using either two-tailed Student*’*s *t* test (for comparison between two groups) or analysis of variance followed by Bonferroni post-hoc test (for comparison between multiple groups). The images with the value closest to the mean value were selected as representative images. A *P* value <0.05 was considered statistically significant.

## Results

### Specific inactivation of Hdac3 in developing epicardium results in ventricular wall hypoplasia

To study the potential role of *Hdac3* in the epicardium during heart development, we specifically ablated *Hdac3* using *Wt1*^*CreERT2* 17^, a tamoxifen inducible epicardial Cre mouse, and *Hdac3* floxed mice^42^. As expected, after tamoxifen induction at E8.5, *Hdac3* was effectively and specifically deleted in the epicardium, whereas its expression in non-epicardial cells, such as cardiomyocytes, was unaffected (Fig. 1A). We did not observe any leakiness of Cre in the *Wt1*^*CreERT2*^ allele (Supplemental Fig. 1). By lineage tracing, we found that *Hdac3* deficient EPDCs were significantly fewer than the littermate control EPDCs at E14.5 (Supplemental Fig. 2). Both epicardium and EPDCs are major contributors to myocardial growth through cell-cell crosstalk^5^. Thus, we assessed the myocardia of epicardial *Hdac3* deficient (*Hdac3*^*eko*^) hearts and found that the ventricular free wall development was markedly affected in *Hdac3*^*eko*^ hearts. The compact layer in *Hdac3*^*eko*^ hearts was significantly thinner than in the littermate control hearts (Fig. 1B), whereas there was no apparent morphological phenotypic difference in the *Hdac3*^*eko*^ trabecular myocardia as compared to the littermate control hearts.

**Figure 1.**
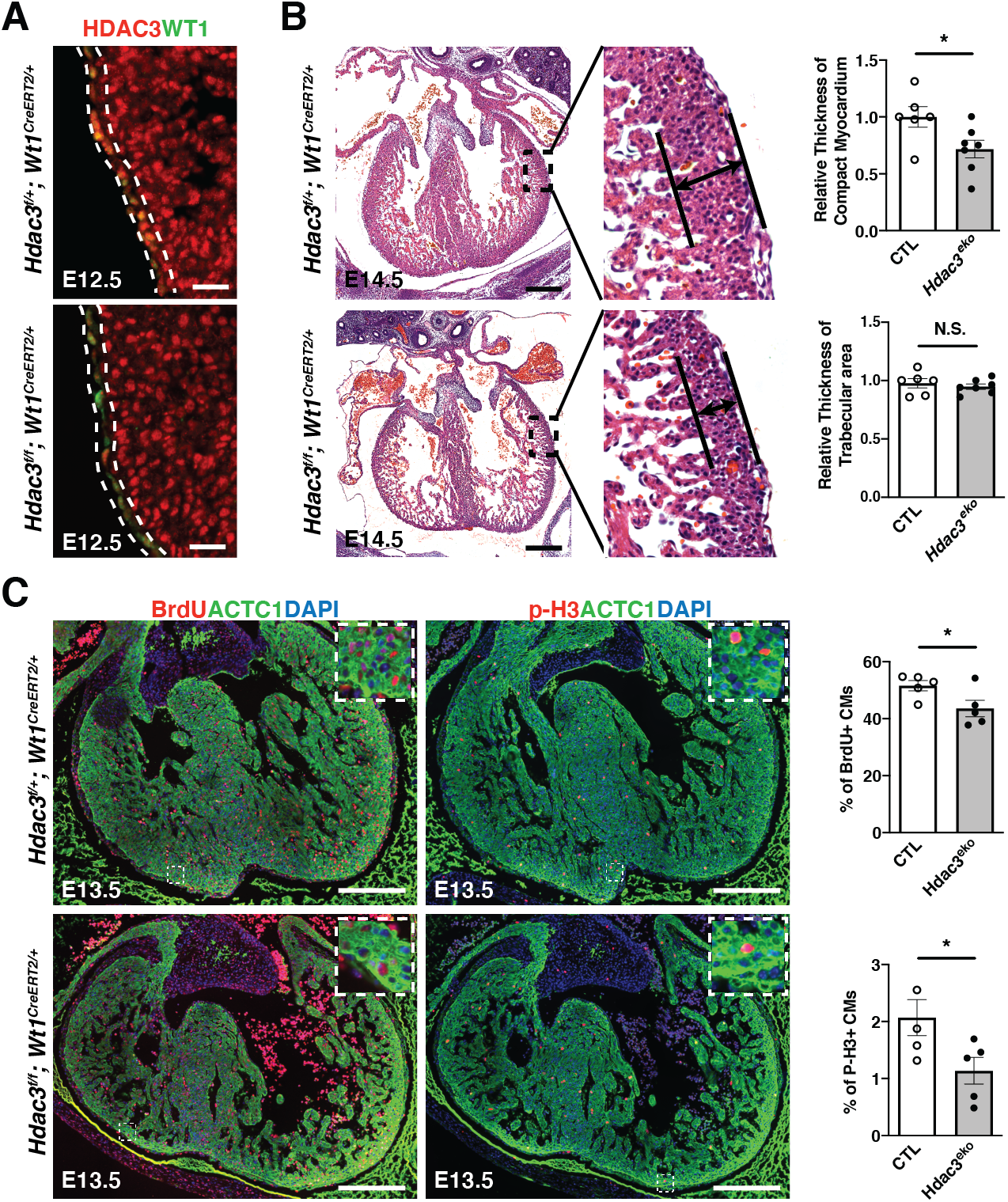
Epicardial deletion of *Hdac3* resulted in hypoplasia of ventricular compact wall. **(A)** Representative immunofluorescence staining of HDAC3 and WT1 in *Hdac3*^*eko*^ (*Hdac3*^*f/f*^; *Wt1*^*CreERT2/+*^) hearts and control (CTL; *Hdac3*^*f/+*^; *Wt1*^*CreERT2/+*^) hearts. Tamoxifen was given to dams intraperitoneally (150 mg/kg body weight) at E8.5 (scale bars, 25 μm). **(B)** Representative H&E staining of *Hdac3*^*eko*^ and CTL hearts. Quantification is shown on the right (\**P*<0.05 by Student*’*s *t*-test; scale bars: 250 μm). **(C)** Representative immunofluorescence staining BrdU and p-H3 in in *Hdac3*^*eko*^ hearts and CTL hearts. Quantification is shown on the right (\**P*<0.05 by Student*’*s *t*-test; scale bars: 250 μm).

Next, we investigated whether altered cell proliferation and/or apoptosis contributes to the ventricular wall hypoplastic phenotypes in *Hdac3*^*eko*^ hearts. At E13.5, we found that the percentage of p-H3+ or BrdU+ myocytes was significantly lower in *Hdac3*^*eko*^ compact myocardia as compared to the littermate control hearts (Fig. 1C). In contrast, there was no significant difference in cell apoptosis between *Hdac3*^*eko*^ hearts and the littermate control hearts (Supplemental Fig. 3). These findings are consistent with the hypoplastic cardiac phenotypes seen in *Hdac3*^*eko*^ hearts (Fig. 1B).

### Reduced expression of FGF9 and IGF2 in Hdac3 deficient epicardial cells contributes to deceased cardiomyocyte proliferation

To further understand the potential molecular mechanisms contributing to the hypoplastic ventricular wall phenotype in *Hdac3*^*eko*^ hearts, we effectively deleted *Hdac3* in immortalized MECs using CRISPR/CAS9 technology (Fig. 2A). *Hdac3* KO MECs grow slower than *Hdac3* EV MECs. By performing bulk RNA sequencing and gene ontology (GO) analysis on *Hdac3* KO and EV MECs, we found 1,681 downregulated genes and 1,549 upregulated genes. GO pathway analyses identified that the top 14 out of 20 significantly affected pathways are all involved in cell division (Fig. 2B). In a search for EPDC secreted growth factors in downregulated genes, we identified *Fgf9* and *Igf2* as our top candidates (volcano plot in Fig. 2C). We further validated the decrease of FGF9 and IGF2 at both mRNA and protein level in *Hdac3* KO MECs (Fig. 2D and E). Consistently, FGF9 and IGF2 were also greatly decreased in *Hdac3*^*eko*^ hearts (Supplemental Fig. 4). These results are consistent with the cardiac growth defects seen in *Hdac3*^*eko*^ hearts.

**Figure 2.**
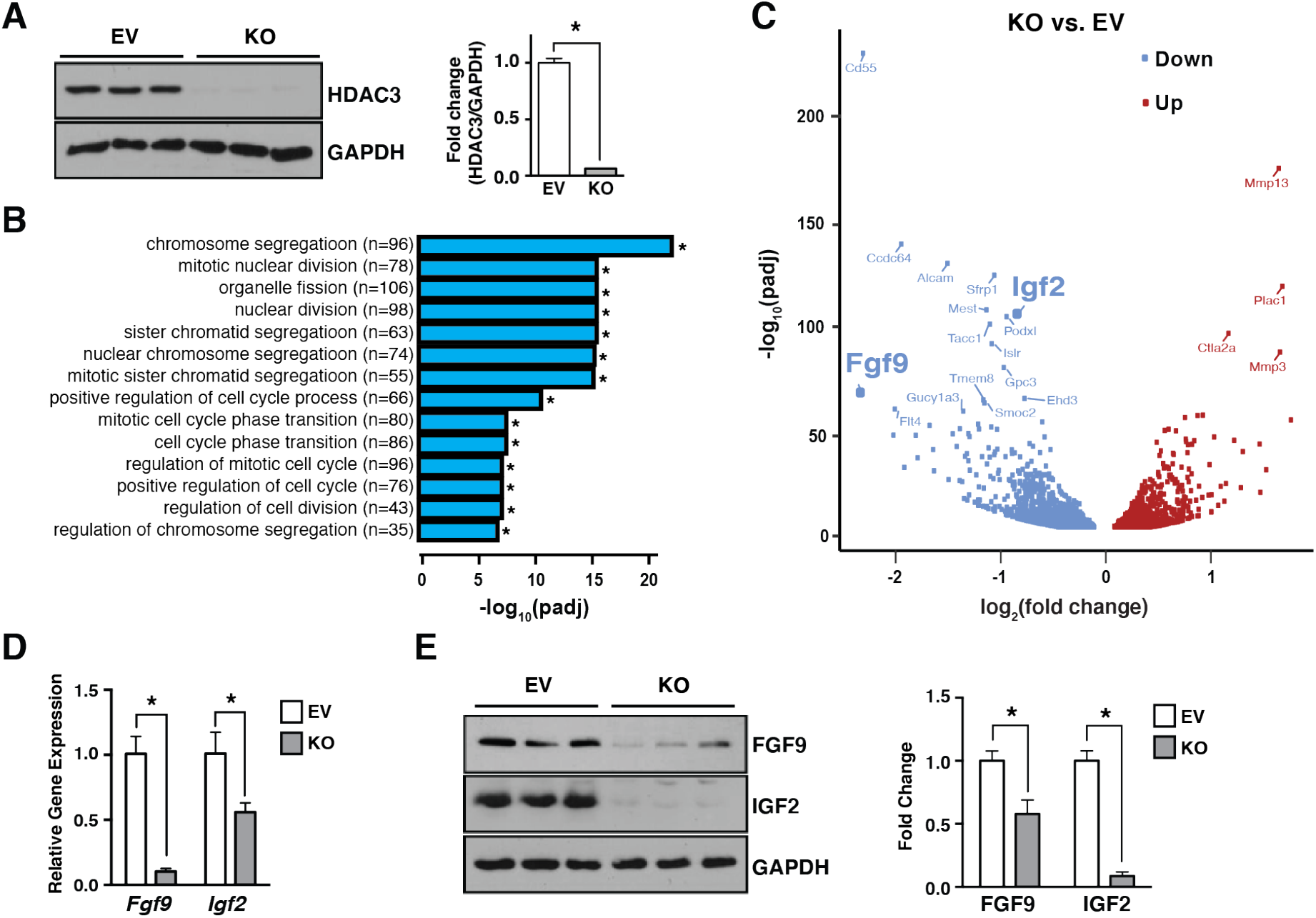
*Hdac3* deletion resulted in downregulation of FGF9 and IGF2. **(A)** Generation of *Hdac3* knockout (KO) and empty vector control (EV) MECs by CRISPR/CAS9. Deletion of *Hdac3* was verified by western blot. **(B)** Gene ontology (GO) pathway analyses and **(C)** volcano plot of RNA Sequencing in *Hdac3* KO and EV MECs. **(D)** Quantification of *Fgf9* and *Igf2* in Hdac3 KO MECs by qRT-PCR. *Gapdh* was used as cDNA loading control (\**P*<0.05 by Student*’*s *t*-test). **(E)** Quantification of FGF9 and IGF2 in *Hdac3* KO and EV MECs by western blot. GAPDH was used as protein loading control (\**P*<0.05 by Student*’*s *t*-test).

Next, we sought to determine whether reduction of FGF9 and IGF2 accounts for decreased cardiomyocyte proliferation. First, we found that FGF9 and IGF2 were significantly decreased in the supernatant from *Hdac3* KO MECs as compared to that from *Hdac3* EV MECs (Fig. 3A). Then, we treated cultured primary embryonic cardiomyocytes isolated from E13.5 *Tnnt2*^*nGFP/+*^ hearts^43^ with supernatants from either *Hdac3* KO MECs or *Hdac3* EV MECs, respectively. *Hdac3* KO supernatant treatment resulted in significant decrease of the percentage of p-H3+ cardiomyocytes and the total number of cardiomyocytes, as compared to *Hdac3* EV supernatant treatment (Fig. 3B and C). Consistently, *Hdac3* KO supernatant attenuated the activation of downstream signaling pathway for cardiomyocyte proliferation, such as p-ERK (Supplemental Fig. 5). Lastly, to determine whether the reduction of FGF9 and IGF2 in *Hdac3* KO supernatants is a major contributor to the decreased cardiomyocyte proliferation, we supplemented *Hdac3* KO supernatant with recombinant mouse FGF9 or IGF2 protein, respectively. Strikingly, supplementation with FGF9 or IGF2 successfully rescued cardiomyocyte proliferation defects by *Hdac3* KO supernatant treatment (Fig. 3B and C). We further confirmed the reactivation of p-FGFR1 or p-IGFR1 by FGF9 or IGF2 supplementation in *Hdac3* KO supernatant treated cardiomyocytes (Supplemental Fig. 5). Altogether, these results suggest that epicardial cells regulate cardiomyocyte growth by inducing the expression of FGF9 and IGF2.

**Figure 3.**
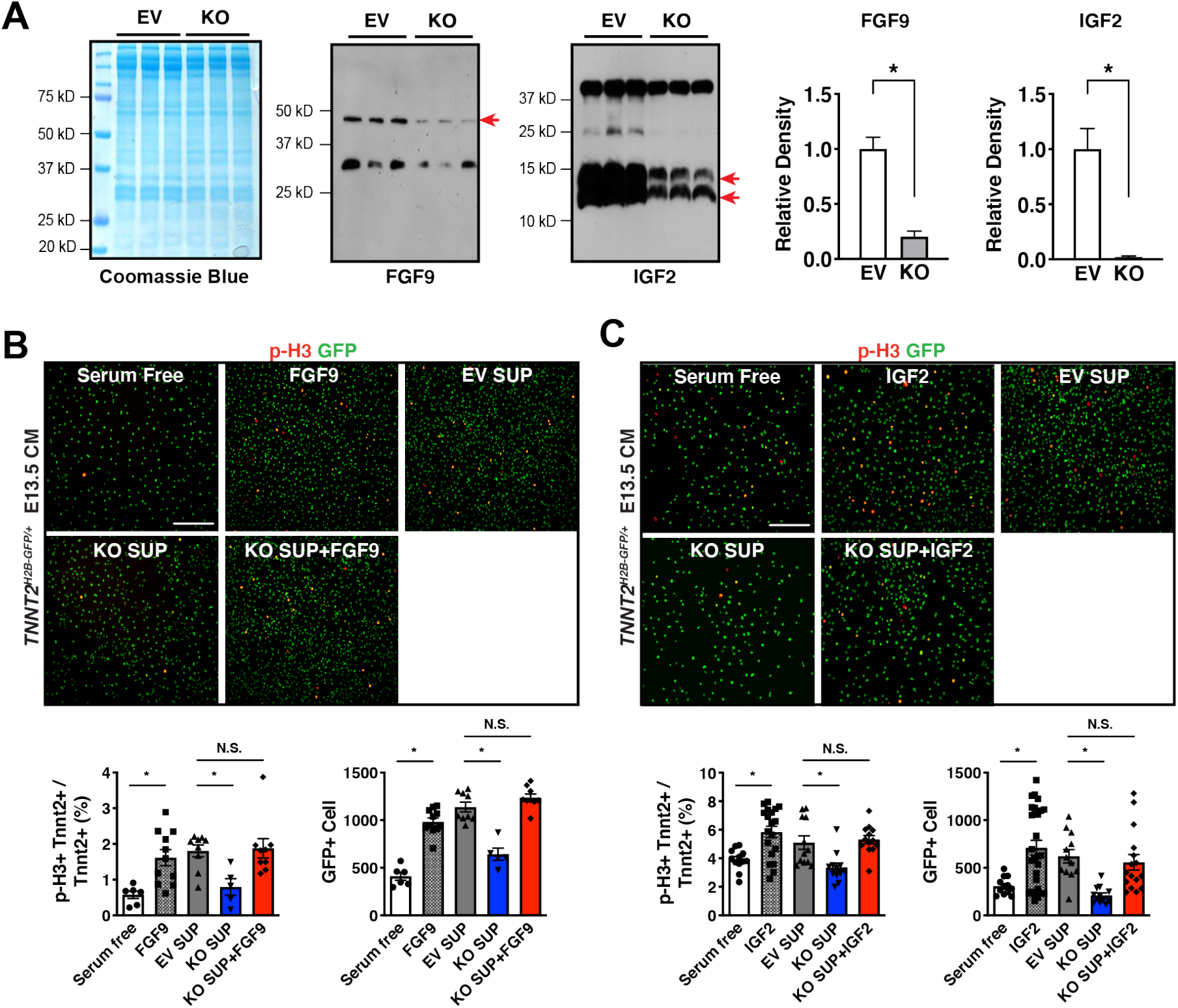
Supplementation of FGF9 or IGF2 rescues cardiomyocyte proliferation defects. **(A)** Secretion of FGF9 and IGF2 from *Hdac3* KO and EV MECs. Coomassie brilliant blue staining of total extracted proteins from supernatants served as protein loading controls. FGF9 and IGF2 in the MEC supernatants were detected by western blot. Arrows point to the target bands. Quantifications are shown on the right. **(B&C)** The effects of MEC supernatants and/or recombinant FGF9 or IGF2 on E13.5 *Tnnt2*^*nGFP/+*^ CM proliferation. Representative immunofluorescence micrographs are shown. Percentage of p-H3+ CMs and total number of CMs were quantitate (\**P*<0.05 by ANOVA followed by Bonferroni post-hoc test; scale bars: 275 μm).

### HDAC3 induces FGF9 and IGF2 expression dependent on its deacetylase enzymatic activities

To determine whether the deacetylase activities are required for HDAC3 to induce the expression of FGF9 and IGF2, we treated MECs with RGFP966, a selective HDAC3 deacetylase inhibitor^52^. Interestingly, RGFP966 treatment significantly decreased FGF9 and IGF2 in a dose-dependent manner (Fig. 4A and B). To further confirm that HDAC3 regulates FGF9 and IGF2 dependent on its enzymatic activities, we performed a set of genetic rescue experiment. As expected, re-expression of lentiviral *Hdac3* in *Hdac3* KO MECs successfully restored the expression of *Fgf9* and *Igf2* (Fig. 4C). In contrast, re-expression of lentiviral *Hdac3* Y298H, a deacetylase dead mutant *Hdac3*^53^, failed to rescue the expression of *Fgf9* and *Igf2* expression (Fig. 4C).

**Figure 4.**
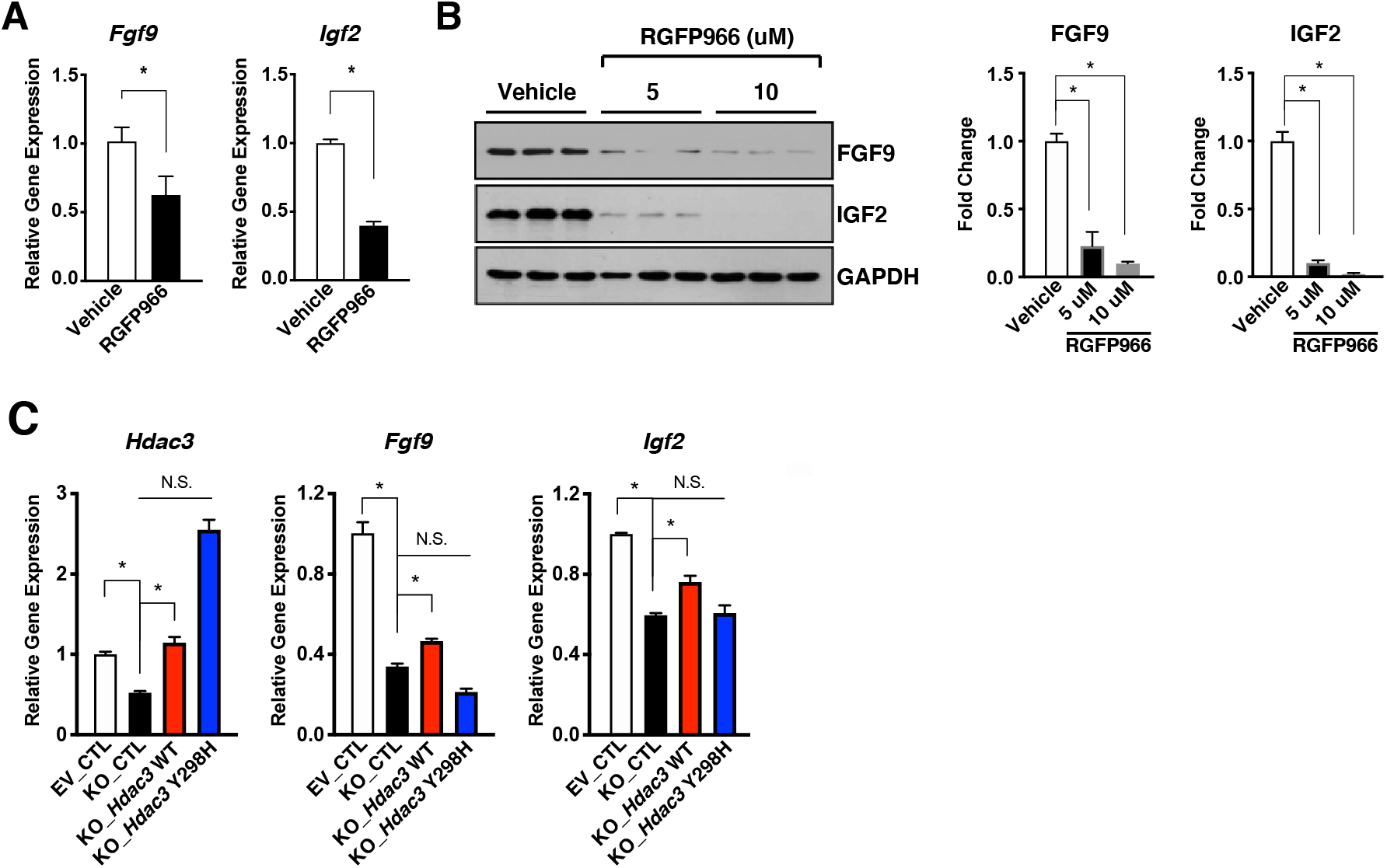
HDAC3 induces the expression of FGF9 and IGF2 dependent on its deacetylase activity. **(A)** Decease of *Fgf9* and *Igf2* mRNAs after RGFP966 (a selective *Hdac3* inhibitor) treatment. MECs were treated with 10 uM RGFP966 or vehicle for 24 hours. mRNA levels were quantified by qRT-PCR. *Gapdh* was used as a cDNA loading control (\**P*<0.05 by Student*’*s *t*-test). **(B)** Decease of FGF9 and IGF2 protein levels after RGFP966 treatment. MECs were treated with 5 or 10 uM RGFP966 or vehicle for 24 hours. FGF9 and IGF2 were quantified by western blot. GAPDH was used as protein loading control (\**P*<0.05 by Student*’*s *t*-test). **(C)** mRNA levels of *Fgf9* and *Igf2* in *Hdac3* KO and EV MECs after 24-hour treatment with *Hdac3* WT, Y298H mutant, or mCherry (CTL) lentivirus. mRNAs were quantified by qRT-PCR (*n*=3, \**P*<0.05 when compared to the CTL group by ANOVA followed by Bonferroni post-hoc test).

### HDAC3 induces expression of FGF9 and IGF2 through repressing miR-322 and miR-503

The expression of FGFs and IGFs can be modulated by miRs^54, 55^, which can be downstream targets of HDACs in certain biological contexts^56^. To identify the potential HDAC3 downstream miR targets, we performed miR sequencing in *Hdac3* KO and EV MECs. Through differential expression analyses, 42 miRs were significantly upregulated in *Hdac3* KO MECs. Among the top 20 hits (fold change equal to or greater than 1.5), we identified 11 miRs (Fig. 5A) that have putative binding sites on either *Fgf9* or *Igf2* using the “DIANA MicroT-CDS” analyses tool^57^. We treated MECs with these 11 miR mimics and found that the treatment of miR-322 mimics or miR-503 mimics significantly inhibited the expression of both *Fgf9* and *Igf2* (Fig. 5B). miR-322 and miR-503 are encoded as one cluster by H19X, which is located in chromosome X, and they share the same “AGCAGC” sequences within the seed region at the 5*’* end. The 3*’* UTRs of both *Fgf9* and *Igf2* harbor putative binding sites for miR-322 and miR-503 (Fig. 5C). We further validated the significant inhibitory effect of miR-322 mimics or miR-503 mimics treatment on the expression of FGF9 and IGF2 by western blot (Fig. 5D). These results indicate that miR-322/miR-503 represses the expression of *Fgf9* and *Igf2*. We further validated the upregulation of miR-322 and miR-503 in *Hdac3* KO MECs (Fig. 5E). Interestingly, miR-322 and miR-503 were similarly significantly upregulated when the deacetylase activity of HDAC3 is inhibited by RGFP966 treatment (Fig. 5E). The significant upregulation of miR-322 and miR-503 was also observed in E13.5 *Hdac3*^*eko*^ hearts as compared to the littermate control hearts (Fig. 5E).

**Figure 5.**
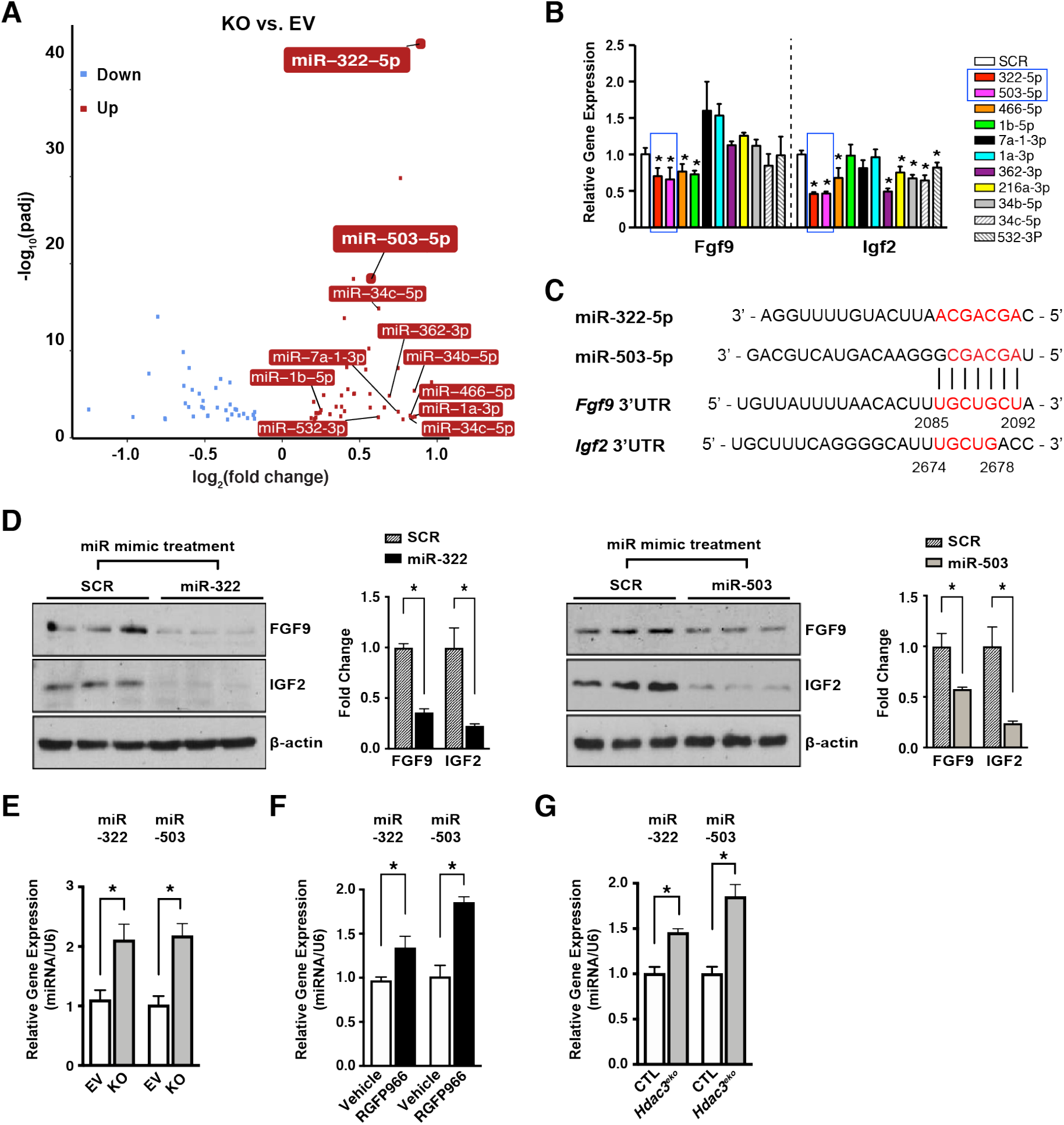
miR-322 and miR-503 repress the expression of FGF9 and IGF2. **(A)** Volcano plot of miR sequencing of *Hdac3* KO and EV MECs. **(B)** Quantification of *Fgf9* and *Igf2* expression in MECs after miR mimics treatment (final concentration:10 nM) by qRT-PCR. *Gapdh* was used as cDNA loading control (\**P*<0.05 by ANOVA followed by Bonferroni post-hoc test). **(C)** miR-322 and miR-503 share high similarity of their seed binding motifs to 3*’*UTRs of *Fgf9* and *Igf2*. Binding motifs or complementary bases are in red. **(D)** Quantification of the expression of FGF9 and IGF2 after miR-322 or miR-503 mimic treatment by western blot. SCR: miR scrambles (\**P*<0.05 by Student*’*s *t*-test). **(E)** Quantification of miR-322 and miR-503 in MECs after miR mimic treatment by qRT-PCR. U6 snRNA was used for normalization (\**P*<0.05 by Student*’*s *t*-test). **(F&G)** Quantification of the expression of miR-322 and miR-503 in *Hdac3* KO MECs, RGFP966-treated MECs, or E13.5 *Hdac3*^*eko*^ hearts by qRT-PCR (\**P*<0.05 by Student*’*s *t*-test).

To determine whether the upregulation of miR-322 and miR-503 has causal effects on the reduction of FGF9 and IGF2 when *Hdac3* is knocked out or inhibited, we knocked down the expression of miR-322 or miR-503 by miRZip lentivirus in *Hdac3* KO MECs (Fig. 6A). Remarkably, knockdown of miR-322 or miR-503 significantly restored the expression of FGF9 and IGF2 in *Hdac3* KO MECs (Fig. 6B). These results suggest that HDAC3 promotes the expression of FGF9 and IGF2 through repressing miR-322/miR-503.

**Figure 6.**
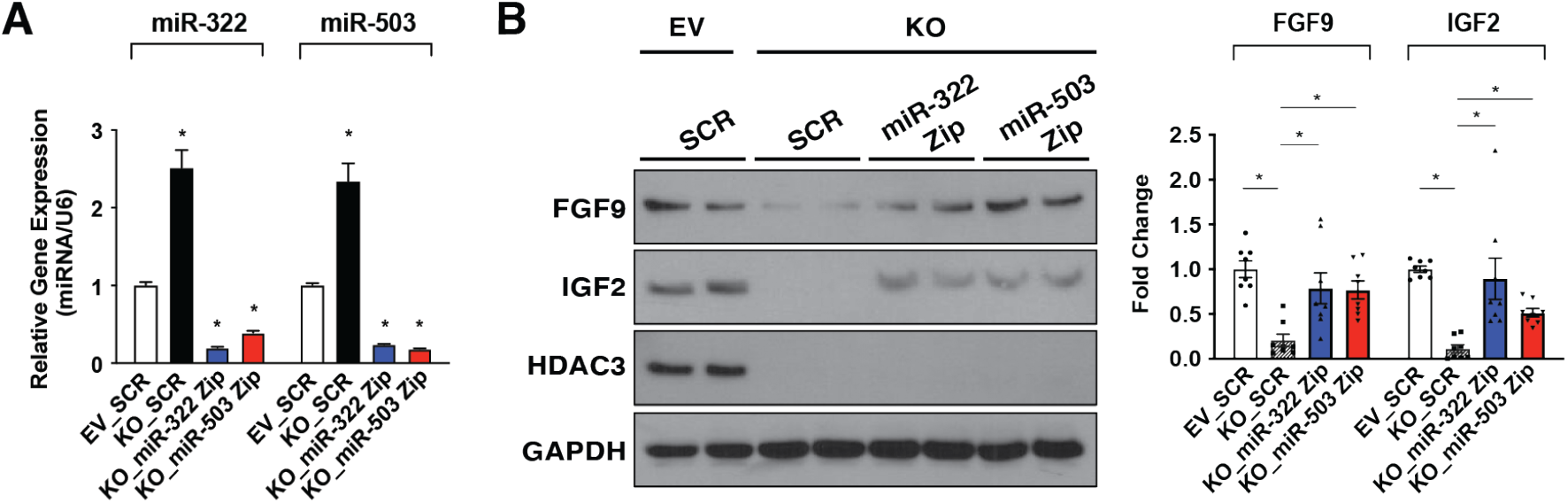
Knockdown of miR-322 or miR-503 restores the expression of FGF9 and IGF2. **(A)** Quantification of miR-322 and miR-503 after *Hdac3* KO or EV MECs were infected with LentimiRa-GFP-miRZip (miR-322 or miR-503) or pGreenPuro Scramble Hairpin control lentivirus (SCR) respectively. miR levels were quantified by qRT-PCR (*n*=3, \**P*<0.05 when compared to the SCR group by ANOVA followed by Bonferroni post-hoc test). **(B)** Quantification of the expression of FGF9 and IGF2 after miRZip lentiviral treatment by western blot (\**P*<0.05 when compared to the SCR group by ANOVA followed by Bonferroni post-hoc test).

### HDAC3 represses miR-322/miR-503 promoter activity

H3K27Ac is a marker for active promoters and a direct downstream target of HDAC3^32^. As expected, *Hdac3* deletion resulted in significant increase of H3K27Ac in MECs (Fig. 7A). Identified by the ENCODE project, the promoter region of miR-322/miR-503 is subject to epigenetic regulations such as H3K27Ac (Fig. 7B). To determine whether *Hdac3* deletion would affect the chromatin accessibility of the miR-322/-503 promoter, we surveyed H3K27Ac binding affinity in *Hdac3* KO MECs by ChIP-qPCR. We found that *Hdac3* deletion significantly increased the binding of H3K27Ac to the miR-322/miR-503 promoter (Fig. 7C), suggesting that *Hdac3* deletion renders the miR-322/miR-503 promoter more accessible to transcriptional factors. To further test whether this increased chromatin accessibility would affect the miR-322/miR-503 promoter activities, we performed an miR-322/miR-503 promoter luciferase reporter assay. We found the luciferase activity was significantly increased in *Hdac3* KO MECs as compared to *Hdac3* EV MECs (Fig. 7D). Strikingly, inhibition of HDAC3 by RGFP966 replicated this result (Fig. 7D), suggesting that the regulation of HDAC3 on the miR-322/miR-503 promoter is dependent on its deacetylase activities.

**Figure 7.**
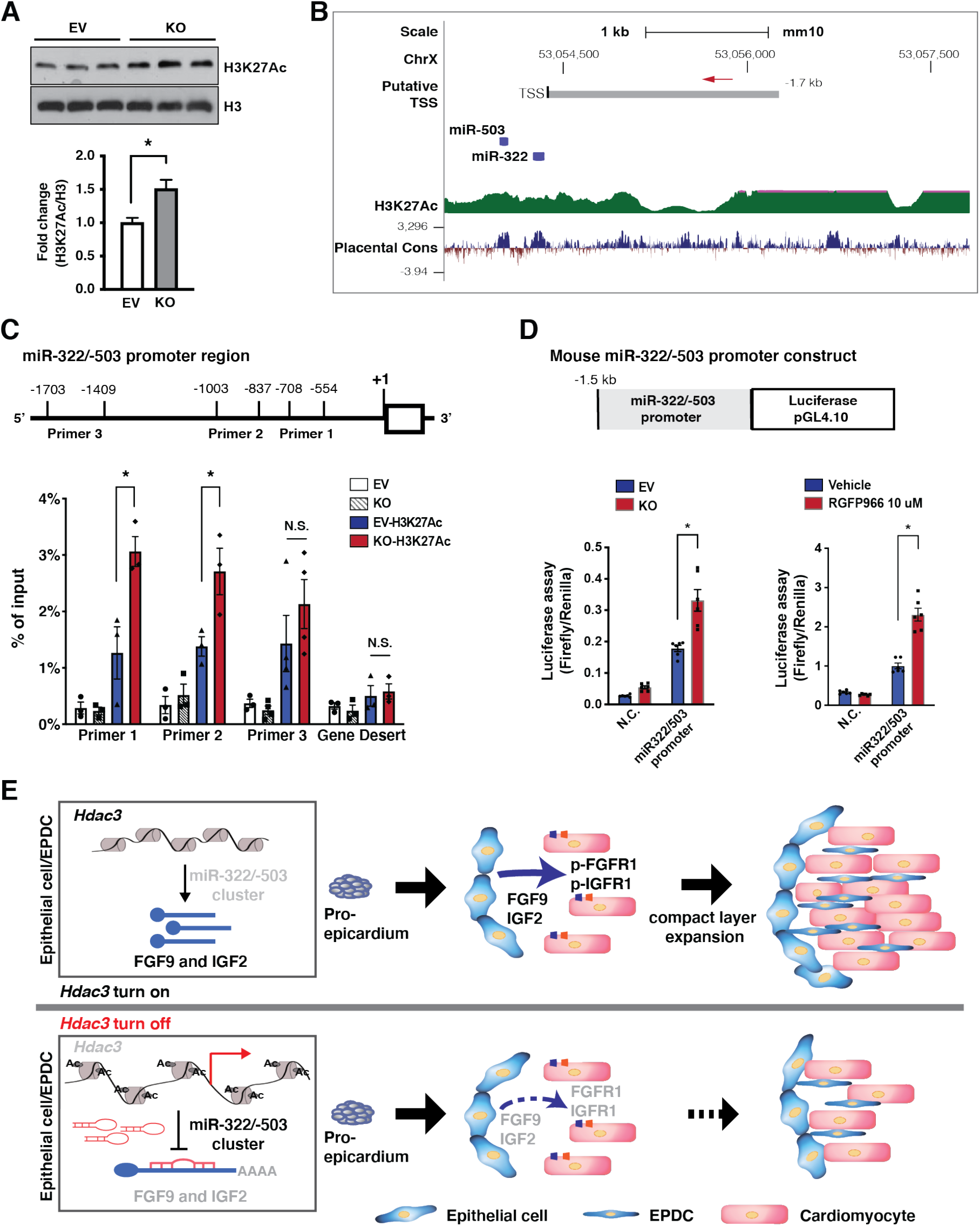
HDAC3 represses miR-322/miR-503 promoter activity. **(A)** Quantification of H3K27Ac in *Hdac3* EV and KO MECs by western blot. **(B)** Schematic diagram of the miR-322/miR-503 locus from UCSC Genome Browser. In the upstream regulatory regions as well as gene bodies, active epigenetic marker H3K27Ac was identified by the ENCODE project. **(C)** Quantification of H3K27Ac binding affinity in the miR-322/miR-503 promoter region in *Hdac3* KO and EV MECs by ChIP-qPCR. Primers targeting a gene desert region were used as a negative control (*n*=3, \**P*<0.05 when compared to the SCR group by ANOVA followed by Bonferroni post-hoc test). **(D)** Dual luciferase reporter assays on the -1.5 kb miR-322/miR-503 promoter when *Hdac3* is either knocked out or inhibited by RGFP966 (final concentration: 10 uM) treatment. The ratio of firefly: Renilla luciferase light units (RLU) was determined 48 hours after transfection (*n*=3, \**P*<0.05 by Student*’*s *t*-test). **(E)** The schematics of the working model. In the developing epicardium, HDAC3 represses the expression of miR-322/miR-503 to release their suppression on the expression of FGF9 and IGF2. When *Hdac3* is deleted, the expression of miR-322/miR-503 is increased, which subsequently suppresses the expression of FGF9 and IGF2 to a stronger extent, and the decrease of FGF9 and IGF2 leads to ventricular wall hypoplasia.

## Discussion

Early elegant studies in avian embryos have revealed that delay or blockade of the epicardium formation leads to decreased cardiomyocyte proliferation in the ventricular free wall without affecting trabecular myocardial development, which is more dependent on the support from the endocardium^4, 58, 59^. Subsequent studies found that epicardium stimulates ventricular wall expansion by providing mitogens to cardiomyocytes such as retinoic acid, FGFs, and IGFs^24, 60, 61^. However, it is unclear how these mitogens are initially induced in the epicardium. Epigenetics has been increasingly recognized as an important regulator of gene transcription in a variety of phycological/pathological processes including cardiac development and congenital diseases^62, 63^. Our current study suggests that HDAC3 induces the expression of FGF9 and IGF2 in the epicardium, and thus stimulates ventricular myocardial wall expansion through paracrine signaling. Our study provides strong evidence that the epigenetics in the epicardium regulates ventricular wall morphogenesis, a new perspective on the mechanisms of cardiac development.

The molecular action of HDAC3 is complex. Nuclear receptor co-repressor 1 (NCoR1 or NCoR) and silencing mediator of retinoic acid and thyroid hormone receptor (SMRT) are both nuclear receptor co-repressors. The NCoR/SMRT complex stoichiometrically recruits HDAC3 and activates its intrinsic deacetylase activity^32^. HDAC3 mainly deacetylates H3K9Ac and H3K27Ac. Deacetylation of these two sites renders chromatin structure unfavorable for the recruitment of transcriptional factors, and thus dampens gene transcription. This is the mechanism by which HDAC3 epigenetically suppresses gene transcription in regulation of skeletal muscle metabolism^64, 65^. However, when HDAC3 determines cardiomyocyte fate, it acts as a recruiter to tether genes to the nuclear periphery for silencing and its deacetylase enzymatic activity is dispensable^66^. Interestingly, several recent studies highlight a critical function for HDAC3 as a gene activator, rather than a gene repressor^66-69^. Our current study demonstrates that in the developing epicardium, HDAC3 works as a gene repressor to suppress the expression of miR-322/miR-503 through its deacetylase activities. As HDAC3 lacks intrinsic DNA binding capacity, it needs to interact with cofactors to modulate promoter/enhancer activities. Whether HDAC3 acts as a repressor or an activator might depend on the nature of its interacting partners and cell context. To identify the potential targets of HDAC3 in the epicardium, we narrowed down to FGF9 and IGF2 by both unbiased screen (RNA-Seq) and candidate (known growth factors) approaches. It is possible that other unaccounted dysregulated genes caused by *Hdac3* deletion might contribute to the hypoplastic ventricular wall phenotype. We will explore those possibilities in future studies.

miR-322 (ortholog of human miR-424) and miR-503 belong to the miR-15/107 family. They are encoded as one cluster by H19X^70^. miR-322 and miR-503 regulate many fundamental processes such as cell proliferation, death, differentiation, metabolism, and stress response. Thus, they are widely implicated in many disorders such as cardiovascular disease, neural disease, and cancer^71, 72^. During heart development, miR-322/miR-503 has been shown to drive early cardiac progenitor cells toward cardiomyocyte lineage, although the underlying mechanism is undetermined^**73**^. Our current study demonstrates that HDAC3 represses miR-322 and miR-503 expression to induce the expression of several major growth ligands including FGF9 and IGF2 for ventricular wall cardiomyocyte proliferation (Fig. 7). This is the first line of study to demonstrate that the myocardial growth cues from the epicardium are tightly controlled by double inhibitory epigenetic regulation with HDACs and microRNAs. HDAC3 modulates the accessibility of miR-322/miR-503 promoter (Fig. 7). At this point, it is unclear what transcription factors interact with HDAC3 to regulate the expression of miR-322/miR-503. We will attempt to identify them in future studies.

Many forms of CHD, such as hypoplastic left heart syndrome, are associated with defective cardiac myogenesis, which further compromises cardiac contractile function^74^. Proper cardiac myogenesis requires precise coordination among multiple cell types in the developing heart, rather than just requiring cardiomyocytes. Nonmyocytes including EPDCs provide key growth signals to the adjacent compact myocardium in a paracrine fashion^4^. Our findings suggest that disrupted epicardial signaling (FGF9 and IGF2) significantly compromises the compact wall expansion in *Hdac3*^*eko*^ hearts. Epicardium also plays important roles during adult heart regeneration and repair^5, 7-9^. In future studies, it will be interesting to determine whether manipulation of this HDAC3*—*miR-322/miR-503*—*FGF9/IGF2 axis affects the outcome of heart repair after injuries.

During heart development, EPDCs mainly give rise to interstitial cardiac fibroblasts and coronary smooth muscle cells^16-19^, maybe to cardiomyocytes and endothelial cells with controversial results^17, 75-78^. We found that the total number of EPDCs is significantly decreased in *Hdac3*^*eko*^ hearts, whereas the percentage of contributions to each cell type was not affected (Supplemental Fig. 2). It is not well understood how the lineage contribution by EPDCs is regulated. Growth factors (e.g., FGF10) can promote this process by either inducing EMT or stimulating the proliferation of epicardium-derived terminal cells^6, 61, 79, 80^. It is possible that the reduction of FGF9 and IGF2 in *Hdac3*^*eko*^ hearts accounts for decreased derivation of epicardial lineages.

In summary, we demonstrated that epicardial HDAC3 orchestrates ventricular wall expansion by inducing FGF9 and IGF2 expression through repressing miR-322/miR-503. Our study provides strong evidence that epigenetic factors such as HDAC3 play pivotal roles for the expression of these paracrine growth signals in the epicardium. Our findings strengthen the importance of epicardial paracrine signaling for myocardial development, which is implicated in the pathogenesis of CHDs and adult heart regeneration.

## Acknowledgments

We gratefully acknowledge Dr. Mitchell Lazar (University of Pennsylvania, Philadelphia, Pennsylvania, USA) for providing *Hdac3*^*flox/+*^ mice. We are also very grateful to Dr. Zheng Sun at Baylor College of Medicine for providing *Hdac3* plasmids. We also thank Dr. Sari D. Holmes at the University of Maryland School of Medicine for her critical reading and editing of the manuscript.

## Disclosure

None.

### Abbreviations

BrdU: 5-bromo-2*’*-deoxyuridine
FBS: fetal bovine serum
HDAC3: histone deacetylase 3
FGF9: fibroblast growth factor 9
IGF2: insulin-like growth factor 2
GAPDH: glyceraldehyde 3-phosphate dehydrogenase
PBS: phosphate-buffered saline
PCR: polymerase chain reaction

