## Supplemental Data for "Epicardial HDAC3 promotes myocardial growth through a novel microRNA pathway"

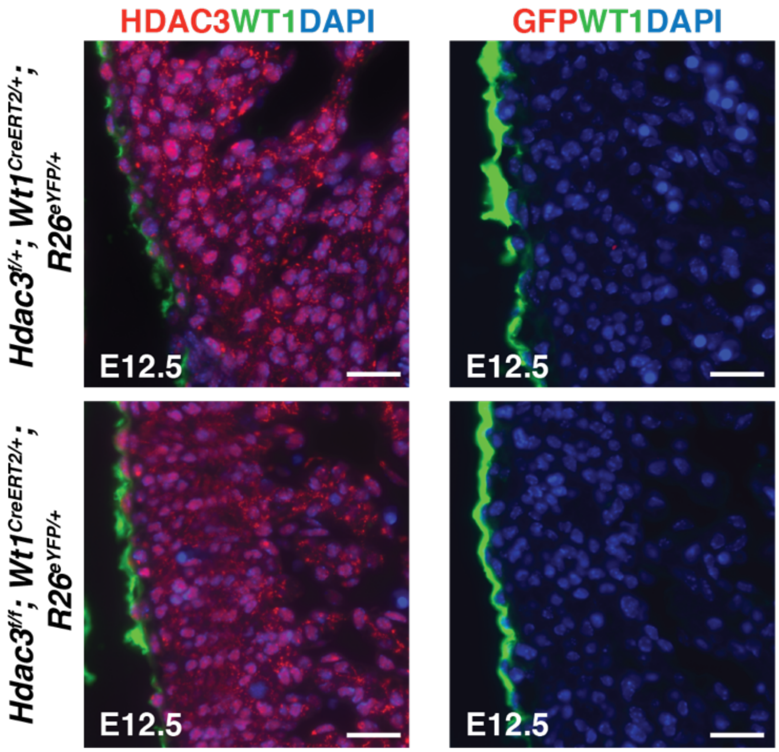


**Supplemental Figure 1. No leakage of *Wt1*$^{CreERT2/+}$.**

Representative micrographs of HDAC3 and GFP immunofluorescence staining of E12.5 *Hdac3*$^{f/+}$*; Wt1*$^{CreERT2/+}$*; R26R*$^{eYFP/+}$ and *Hdac3*$^{f/f}$*; Wt1*$^{CreERT2/+}$*; R26R*$^{eYFP/+}$ hearts. Corn oil was given to dams intraperitoneally (150 mg/kg body weight) at E8.5 (scale bars: 25 μm). eYFP immunosignal was detected by GFP antibody. In the absence of tamoxifen administration, there was neither *Hdac3* deletion nor eYFP reporter activity in the epicardium.

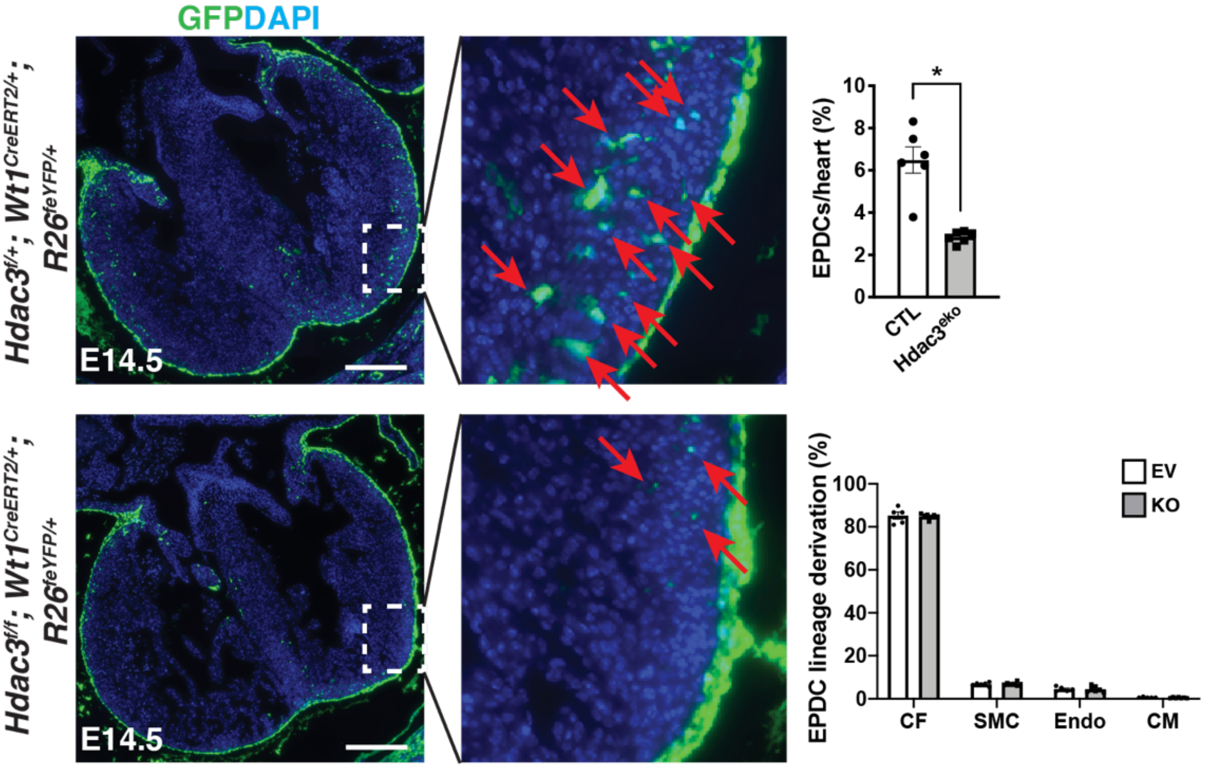


**Supplemental Figure 2. Reduction of EPDCs in *Hdac3$^{eko}$* hearts.**

Representative micrographs of GFP immunofluorescence staining of E14.5 *Hdac3$^{f/+}$; Wt1$^{CreERT2/+}$; R26R$^{eYFP/+}$* (control [CTL]) and *Hdac3$^{f/f}$; Wt1$^{CreERT2/+}$; R26R$^{eYFP/+}$* (*Hdac3$^{eko}$*) hearts. Quantifications of percentage of EPDCs/heart and derivation percentage of each cell type are shown on the right (**P*<0.05 by Student's *t*-test; CF, cardiac fibroblast (Vimentin+); SMC, smooth muscle cells (smMHC11+); Endo, endothelial cells (CD31+); CM, cardiomyocyte (ACTC1+); scale bar, 250 µm). EPDCs (GFP+, denoted by red arrows) were significantly fewer in *Hdac3$^{eko}$* hearts as compared to CTL hearts, whereas the contribution to each lineage by EPDCs was not significantly different between *Hdac3$^{eko}$* and CTL hearts.

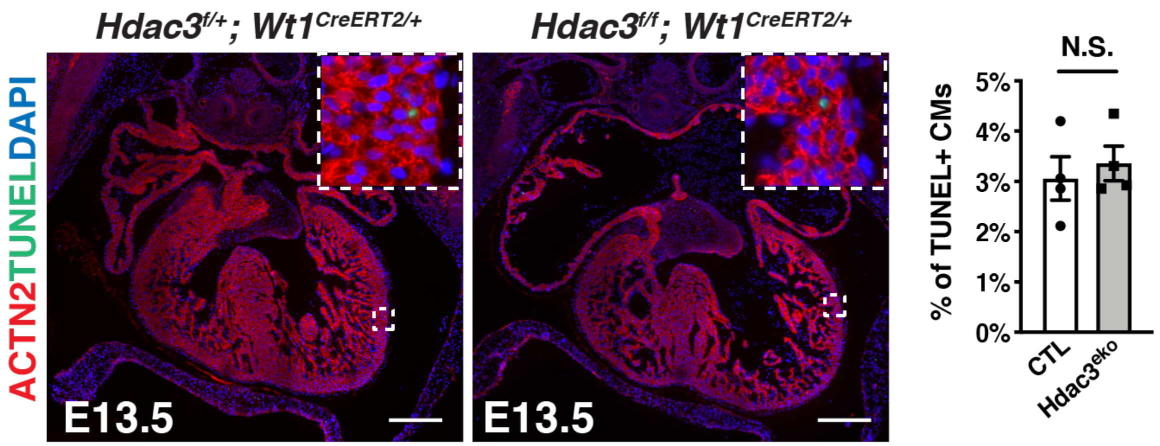


**Supplemental Figure 3. No significant change for cell death in *Hdac3$^{eko}$* hearts.**

Representative micrographs of TUNEL staining of E13.5 *Hdac3$^{f/f}$; Wt1$^{CreERT2/+}$*(*Hdac3$^{eko}$*) and *Hdac3$^{f/+}$; Wt1$^{CreERT2/+}$*(CTL) and hearts. TUNEL+ signals are in green. Quantification of TUNEL+ cardiomyocytes (CMs) is shown on the right (N.S., not significant; scale bars: 250 μm).

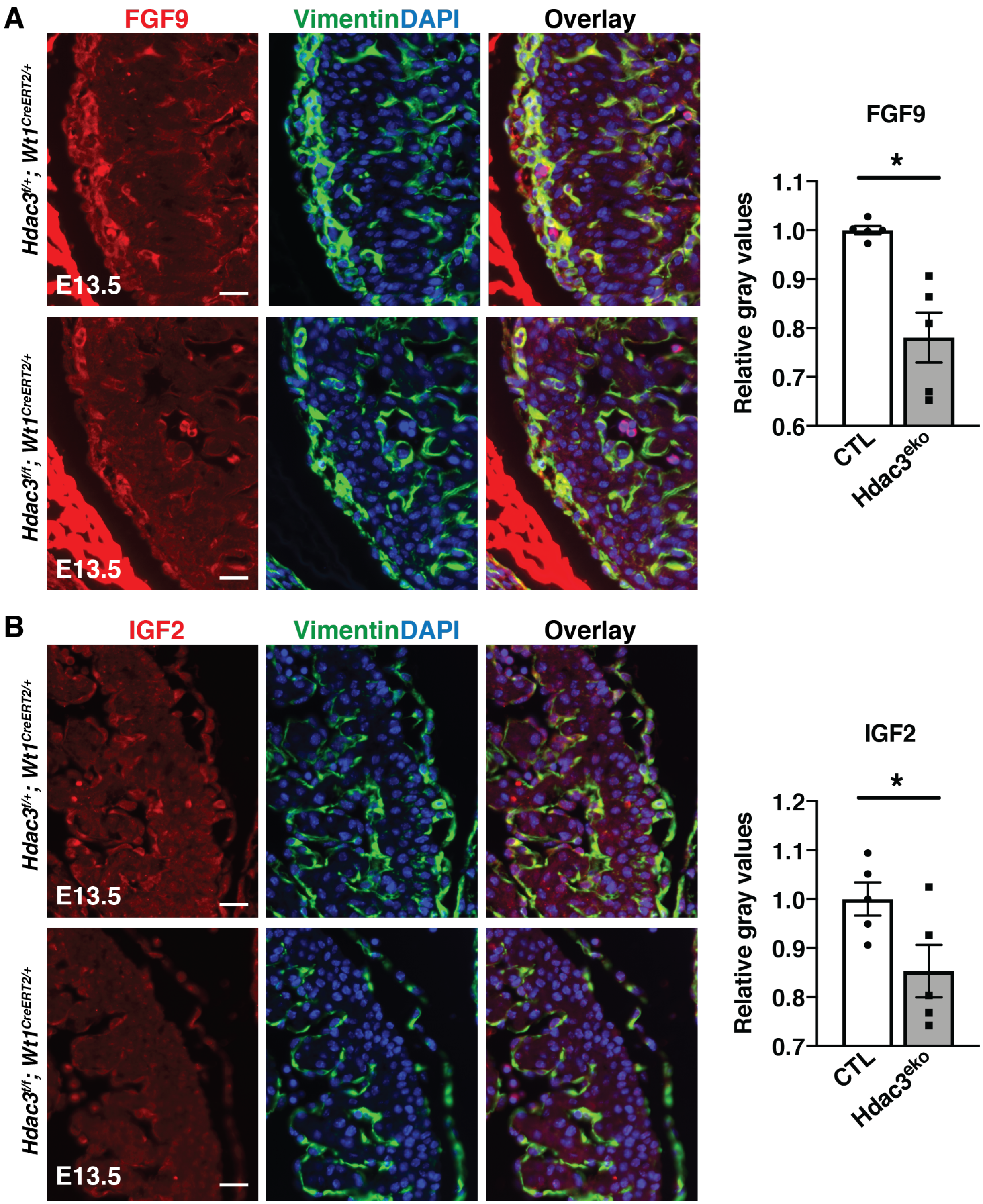

A
FGF9
VimentinDAPI
Overlay
Hdac3f/+; Wt1CreERT2/+
Hdac3f/f; Wt1CreERT2/+
E13.5
FGF9
*
Relative gray values
1.1
1.0
0.9
0.8
0.7
0.6
CTL
Hdac3eko
B
IGF2
VimentinDAPI
Overlay
Hdac3f/+; Wt1CreERT2/+
Hdac3f/f; Wt1CreERT2/+
E13.5
IGF2
*
Relative gray values
1.2
1.1
1.0
0.9
0.8
0.7
CTL
Hdac3eko

**Supplemental Figure 4. Reduced expression of FGF9 and IGF2 in *Hdac3*$^{eko}$ hearts.**

Representative immunofluorescence staining of FGF9 **(A)** and IGF2 **(B)** on E13.5 *Hdac3*$^{f/f}$*; Wt1*$^{CreERT2/+}$*; R26R*$^{eYFP/+}$ and (*Hdac3*$^{eko}$) *Hdac3*$^{f/+}$*; Wt1*$^{CreERT2/+}$*; R26R*$^{eYFP/+}$ (CTL) hearts. Vimentin was used to mark cardiac fibroblasts, cardiac endotheliccal cells and the epicardium. Quantifications of immunofluorescence intensity of FGF9 and IGF2 are shown on the right (scale bars: 25 μm).

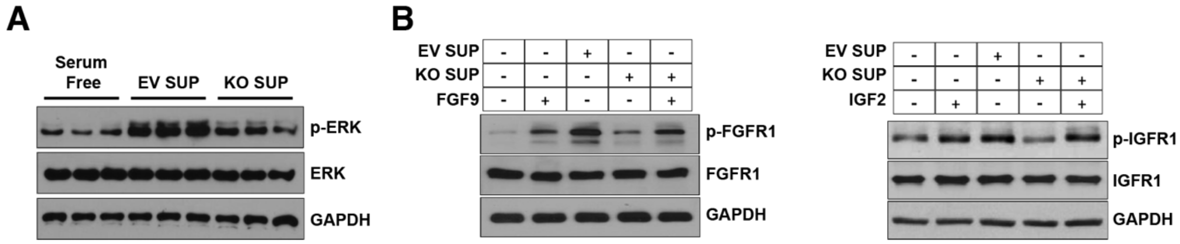


**Supplemental Figure 5. The downstream signaling of FGF9 or iGF2 in cultured cardiomyocytes.** Representative western blots of p-ERK, p-FGFR1, or p-IGF1R in serum-starved cultured E13.5 cardiomyocytes after treatment of MEC supernatants and/or mouse recombinant FGF9 or IGF2 proteins (final concentration: 100 ng/ml).

### Full unedited gels for

**Figure 2A**

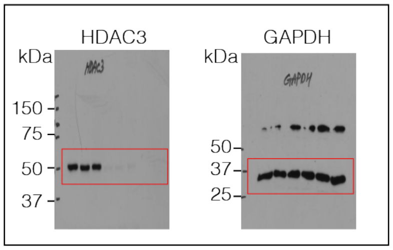


**Figure 2E**

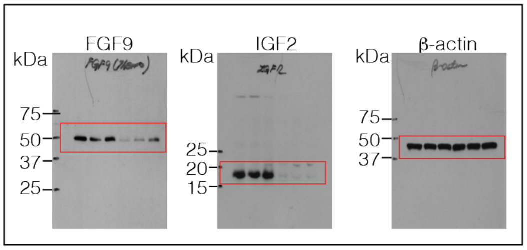


**Figure 4B**

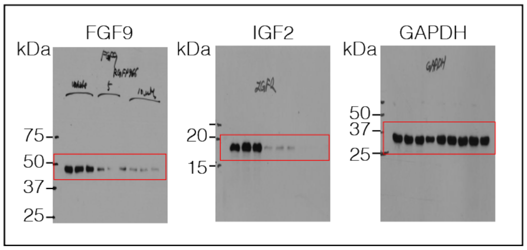


**Figure 5D**

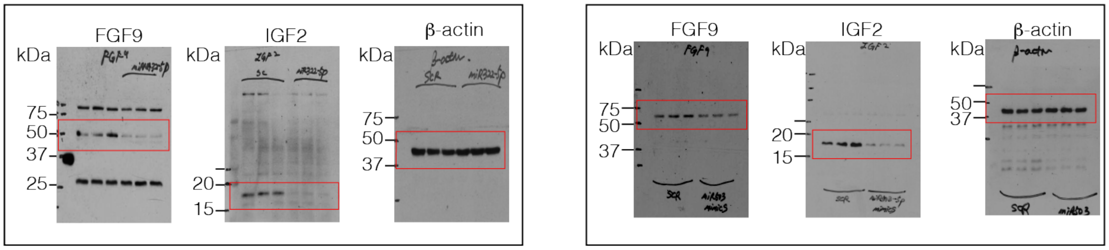


**Figure 6B**

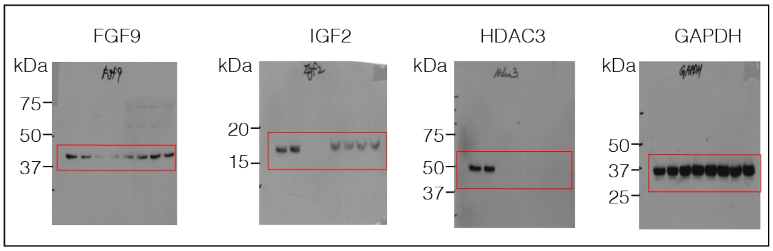


**Figure 7A**

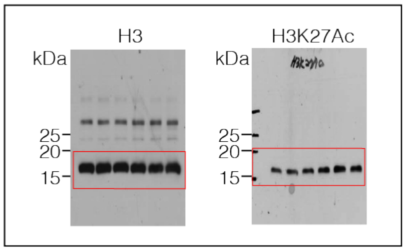


**Supplemental Figure 6. Documentation of full scans of Western blots.**

**Supplemental Table 1. qRT-PCR primers**

| | Forward | Reverse |
|---|---|---|
| ***Fgf9*** | 5’-GGGGAGCTGTATGGATCAGA-3’ | 5’-TCCCGTCCTTATTTAATGCAA-3’ |
| ***Igf2*** | 5’-CGCTTCAGTTTGTCTGTTCG-3’ | 5’-GCAGCACTCTTCCACGATG-3’ |
| ***Gapdh*** | 5’-TCCTGGTATGACAATGAATACGGC-3’ | 5’-TCTTGCTCAGTGTCCTTGCTGG-3’ |

**Supplemental Table 2. ChIP qRT-PCR primers**

| | Forward | Reverse |
|---|---|---|
| **Primer 1** | 5’-GGATGGTTTTTGTGCTTTCC-3’ | 5’-TAAGCCACGCCA CTGAAAAT-3’ |
| **Primer 2** | 5’-CAACTTAAGGAGTGGGGCTGT-3’ | 5’-CAATGAATGCTGGGTCCTTT-3’ |
| **Primer 3** | 5’-GCATGGCATCTGCAACATTA-3’ | 5’-CTCACTCCCTGGGTTTGTGT-3’ |
| **Gene Desert** | 5’-CAGCATGAAAATGGAGGTCA-3’ | 5’-TGAGGGTAAAGGTGCTTGCT-3’ |

**Supplemental Table 3. Antibody used for immunofluorescence or western blot**

| Antibody | Species | Vendor | Catalog # |
|---|---|---|---|
| **BrdU** | Mouse | eBioscience | 14-5071-80 |
| **p-H3** | Rabbit | Cell Signaling | 9701S |
| **ACTC1** | mouse | ARP | 03-61075 |
| **ACTC1** | Rabbit | Abcam | Ab46805 |
| **WT1** | Mouse | Santa Cruz | sc-7385 |
| **GFP** | Goat | Abcam | ab6673 |
| **HDAC3** | Rabbit | Abcam | ab7030 |
| **HDAC3** | Rabbit | Santa Cruz | Sc-11417 |
| **CD31** | Rat | Dianova | Dia-310 |
| **smMHC11** | Mouse | Abcam | Ab683 |
| **Vimentin** | Rabbit | Cell Signaling | 5741 |
| **IGF2** | Goat | Thermo Fisher | PA5-47946 |
| **FGF9** | Rabbit | Abcam | ab206408 |
| **FGF9** | Mouse | Santa Cruz | Sc-8413 |
| **FGFR1** | Rabbit | Cell Signaling | 9740S |
| **p-FGFR1 (Tyr653/654)** | Rabbit | Cell Signaling | 52598S |
| **IGFR1** | Rabbit | Cell Signaling | 3027S |
| **p-IGFR1** | Rabbit | Sigma | SAB4300069 |
| **ERK** | Mouse | BD Biosciences | 610123 |
| **p-ERK** | Rabbit | Cell Signaling | 9101 |
| **H3K27Ac** | Rabbit | Abcam | ab4729 |
| **H3** | Rabbit | Abcam | ab176842 |
| **ß-actin** | Rabbit | Cell Signaling | 4970 |
| **GAPDH** | Mouse | Proteintech | HRP-60004 |
